# Mig6 decreases hepatic EGFR activation and survival during saturated fatty acid-induced endoplasmic reticulum stress

**DOI:** 10.1101/2020.11.24.380527

**Authors:** Andrew J. Lutkewitte, Yi-Chun Chen, Jeffrey L. Hansen, Patrick T. Fueger

**Affiliations:** Department of Cellular and Integrative Physiology, Indiana University School of Medicine, Indianapolis, IN 46202; Department of Pediatrics, Indiana University School of Medicine, Indianapolis, IN 46202; Herman B. Wells Center for Pediatric Research, Indiana University School of Medicine, Indianapolis, IN 46202; Department of Molecular and Cellular Endocrinology, Beckman Research Institute of the City of Hope, Duarte, CA 91010

**Keywords:** ER stress, hyperlipidemia, apoptosis, liver

## Abstract

Hyperlipidemia associated with obesity and type 2 diabetes (T2D) promotes excess hepatic lipid storage (steatosis) and endoplasmic reticulum (ER) stress, thereby reducing hepatic cell proliferation and survival. An important receptor tyrosine kinase controlling liver proliferation and survival is the epidermal growth factor receptor (EGFR). EGFR expression and activation are decreased during steatosis in humans and several animal models of obesity. Therefore, restoring EGFR activation in obesity-induced ER stress and diabetes could restore the liver’s capacity for survival and regeneration. As an inducible feedback inhibitor of EGFR activity, mitogen-inducible gene 6 (Mig6) is a novel target for enhancing EGFR signaling during diet-induced obesity (DIO) and T2D. Thus, we hypothesized hepatic ER stress induces Mig6 expression and decreases EGFR activation during DIO and diabetes. We identified that Mig6 expression was increased during obesity-induced insulin resistance in C57Bl/6J mice fed a high fat diet. We also discovered that both pharmacological- and fatty acid-driven ER stress increased Mig6 expression and decreased EGF-mediated EGFR activation in primary rat hepatocytes and cell lines. Furthermore, siRNA-mediated Mig6 knockdown restored EGFR signaling and reduced caspase 3/7 activation during ER stress. Therefore, we conclude Mig6 is increased during ER stress in DIO, thereby reducing EGFR activation and enhancing cell death. The implications are the induction of Mig6 during DIO and diabetes may decrease hepatocyte survival, thus hindering cellular repair and regenerative mechanisms.

---

During obesity, insulin resistance creates an extracellular milieu providing ample free-fatty acids for uptake and storage by the liver. The resultant increased hepatic steatosis provokes a variety of pathophysiological consequences including hepatic endoplasmic reticulum (ER) stress, exacerbated insulin resistance, and even cell death (33, 43). ER stress negatively impacts a variety of cellular processes, including cell proliferation and survival (41). Several signaling cascades emanating from receptor tyrosine kinases (RTKs) regulate cell proliferation and survival, and these pathways may become compromised during conditions of stress (1, 18, 41). In hepatocytes for example, a major RTK governing cell proliferation and survival is the epidermal growth factor receptor (EGFR). EGFR is activated by several ligands including EGF, TGFα, amphiregulin, and HB-EGF (3, 19, 23, 32, 34). Upon activation, EGFR initiates proliferation by inducing cyclins and suppressing cyclin dependent kinase inhibitors, which together promote the G1-S phase transition (9). Conversely, inhibiting EGFR activation increases cellular caspase 3/7 activity, decreases viability, and reduces expression of prosurvival factors including bcl-2 and survivin (35, 38). Thus, EGFR signaling can control hepatocyte proliferation and the balance between cell survival and death through multiple mechanisms.

Under normal conditions, total hepatic mass is tightly protected; when a loss of hepatic tissue occurs, the liver (and other organs) responds by initiating programs (e.g., EGFR signaling) aimed at increasing the numbers of hepatocytes. For example, the secretion and circulating concentration of many EGFR ligands and subsequent hepatic EGFR activation increases following partial hepatectomy in mice (3, 16, 19, 21, 23, 28, 49). Moreover, studies using liver-specific EGFR ablation demonstrate EGFR is required for complete liver regeneration and stress response kinase, p38, activation following partial hepatectomy (32). Additionally, shRNA-mediated knockdown of EGFR in hepatectomized rat liver reduced BrdU labeling and gene expression of several cyclins. Alternatively, these mice exhibit increased expression of pro-apoptotic genes such as BID, Bak1, and caspases 3/7 and reduced expression of the pro-survival gene surviving, again highlighting the coupling of proliferation and survival controlled by EGFR signaling (35).

Pathologically, hepatic EGFR signaling is decreased in mouse models of obesity, and following partial hepatectomy, the regenerative response in these mice is impaired (10). Importantly, in both mice and human liver tissue, EGFR expression has been negatively correlated with steatosis. Steatosis itself negatively affects postsurgical outcomes in human patients following partial hepatectomy and is associated with increased hepatic apoptosis (10, 27, 43). Thus, understanding the synergistic response of decreased proliferation and survival in liver cells of patients faced with obesity and identifying therapeutic targets is essential for improving overall hepatic health and function. Unfortunately though, the mechanisms for obesity-impaired regeneration and survival as well as hepatic EGFR dysfunction are not well known. Therefore, we set out to define a possible mechanism through which the ER stress-inducible feedback inhibitor of EGFR activation, mitogen inducible gene 6 (Mig6), decreases hepatic EGFR activation during pharmacological-, fatty acid-, and obesity-induced ER stress.

Physiologically, EGFR signaling activates the transcription of several inducible feedback inhibitors, including Socs4, Socs5, Lrig1, and Mig6, which serve to shut off EGFR activation and signaling in a classical negative feedback manner (39). However, Mig6 is not only induced by EGFR activation but also upregulated by a cadre of cellular stressors leading to apoptosis in a variety of cell types (5, 6, 26, 36, 45, 47). Inversely, reducing Mig6 with either genetic ablation or siRNA-mediated suppression enhances EGFR activation and mitogen signaling in hepatocytes (36). Additionally, hepatocytes lacking Mig6 exhibit increased EGFR signaling *in vitro* and enhanced liver regeneration following partial hepatectomy *in vivo* (36). Therefore, we hypothesized that obesity-driven ER stress increases Mig6, which correspondingly decreases EGFR activation. Thus, we speculated that reducing Mig6 during ER stress would restore EGFR activation and reduce cell death.

## MATERIALS AND METHODS

### Animal studies

All animals were handled and cared for according to protocols approved by the Indiana University School of Medicine Institutional Animal Care and Use Committee. Mice were maintained on a standard 12-h light-dark cycle and had free access to chow diet and water. For diet studies, six-week-old male C57Bl/6J mice were fed either a low fat diet (LFD, 15 % kcal from fat, breeder chow) or a high fat diet (HFD, 60% kcal from fat, no. D12492, Research Diets) for 15 weeks. Blood was collected from tail veins, and blood glucose was measured using an AlphaTRAK glucometer (Abbott Laboratories, Abbott Park, IL). Insulin was measured in serum from 6 h-fasted mice using an ELISA (Crystal Chem, Downers Grove, IL). For protein studies livers were immediately harvested, weighed, and flash frozen in liquid nitrogen. Frozen liver samples (100 mg) were placed in lysis buffer (1 mL) containing 2% SDS and homogenized (PolyTron PT2100, Kinematica AG, Luzern, Switzerland). Samples were centrifuged, the aqueous layer was removed, and samples were centrifuged again prior to the BCA assay as described below.

### Cell culture

Cell lines were purchased directly from ATCC (Manassas, VA). HepG2 human male hepatocellular carcinoma cells (HB-8065, ATCC) were cultured in DMEM containing 25 mM glucose supplemented with 10% fetal bovine serum (FBS; Sigma Aldrich, St. Louis, MO), 1 mM sodium pyruvate, and penicillin-streptomycin (100 U/mL). H4IIE rat hepatoma (CRL-1548, ATCC) were cultured similar to HepG2 cells with the exception of the DMEM contained 5.5 mM glucose. Primary hepatocytes were isolated from Sprague Dawley rats at Triangle Research Labs (Durham, NC) and shipped the following day; upon arrival, shipping media was replaced with DMEM containing 25 mM glucose and supplemented as above (both dexamethasone and insulin are known to induce Mig6 gene expression and were therefore omitted during palmitate treatments (8, 11)).

### Ligand stimulation and drug treatments

Twenty-four hours prior to ligand stimulation and during stress treatments, FBS-containing media was replaced with media containing 0.1% bovine serum albumin (BSA, Sigma Aldrich). Cells at 70-80% confluency were treated with thapsigargin (Sigma Aldrich), tunicamycin (Sigma Aldrich), or vehicle (DMSO) as indicated. Cells were treated at dose and time indicated with palmitate or vehicle (BSA) using sodium palmitate (Sigma Aldrich) conjugated to BSA in ethanol at 65°C. Cells were stimulated with 50 ng/mL recombinant rat EGF (R&D systems, Minneapolis, MN) for times indicated.

### Western blot analysis and antibodies

HepG2, H4IIE, and primary rat hepatocytes were washed twice with ice cold PBS and lysed in IGEPAL lysis buffer (10% IGEPAL, 10% glycerol, 1 mM NaCl, 1 mM HEPES, and *O*-octyl glucoside, phosphatase and protease inhibitor tablets (Roche, Mannheim, Germany). Samples were vortexed and centrifuged at >20,000 x *g* for 10 min at 4°C. SDS (2%) was added to lysates from cells treated with palmitate, then sonicated and spun at room temperature. The middle aqueous layer was removed and re-centrifuged. Protein content was determined by bicinchoninic acid assay. Protein (30-40 μg) was reduced in LDS loading buffer plus reducing agent (Invitrogen) and boiled at 100°C for 6 min. Samples were loaded onto 10% Bolt Bis-Tris gels (Invitrogen) and separated at 140 volts for 50 min. Proteins were transferred to an Immunobilon-FL Transfer Membrane (Millipore, Bedford, MA) at 110 volts for 50 min in ice cold transfer buffer. Membranes were blocked in Li-Cor Odyssey blocking buffer for 1 h at room temperature and incubated with primary antibody overnight. Membranes were then washed 3 times in TBS and incubated with IRDye 800 or 700 flurophore-labeled secondary antibodies for 1 h. Antibodies were diluted in either 1% w/v polyvinylpyrrolidone or signal enhancer (Nacalai, San Diego, CA) and are listed in Table 1. Protein bands were visualized on the Odyssey System (LI-COR) and quantified using ImageJ software (National Institutes of Health). Boxes around the representative images indicate antibodies probed on the same membrane.

**Table 1.** List of antibodies used

| Antibody | Manufacturer | Product Number | Dilution |
| --- | --- | --- | --- |
| Mig6 | Santa Cruz Biotechnology Inc. | 137154 | 1:500 |
| P-eif2α | Cell signaling | 3398 | 1:1000 |
| eif2α | Cell signaling | 5324 | 1:1000 |
| P-ERK 1/2 | Cell signaling | 9101S | 1:1000 |
| ERK 1/2 | Cell signaling | 4696S | 1:1000 |
| P-EGFR | Cell signaling | 3777S | 1:1000 |
| EGFR | Cell signaling | 4267S | 1:1000 |
| Caspase 3 | Cell signaling | 9662 | 1:1000 |
| JNK | Cell signaling | 9252S | 1:1000 |
| P-JNK | Cell signaling | 9251S | 1:1000 |
| Actin | MP biomedical | 691002 | 1:10000 |
| GAPDH | abcam | 9484 | 1:500 |
| γ-tubulin | Sigma Aldrich (mouse) | T6557 | 1:1000 |
| EGFR | Sigma Aldrich (mouse) | E3138 | 1:1000 |
| FLAG | Sigma Aldrich (mouse) | F3165 | 1:1000 |

### Caspase 3/7 activity assay

HepG2 cells were transfected as before and the following day trypsinized and plated at 40,000 cells per well in black-walled 96-well plates. The following day media was removed and replaced with serum free media containing 0.1% BSA and treated with vehicle (BSA) or palmitate for 8 h. Following palmitate treatment, caspase 3/7 activity was measured using ApoLive-Glo Multiplex Assay (Promega, Madison, WI). Briefly, cells were incubated with Caspase-Glo 3/7 reagent for 30 min and luminescence was measured using a SpectrMax M5 (Molecular Devices, Sunnyvale, CA). Relative luminescence units were normalized to siCon untreated cells and are given as fold increase in caspase 3/7 activity.

### Reverse transcription and quantitative PCR

For RNA isolations, cells were lysed in RLT buffer containing 1% β-mercaptoethanol. Genomic DNA was removed via gDNA column (Qiagen, Valencia, CA), and RNA was isolated with an RNeasy mini prep kit (Qiagen). Complementary DNA was generated using high capacity cDNA kit (Applied Biosystems, Foster City, CA). Quantitative PCR was performed using Taqman Master Mix and Taqman probes as listed in Table 2. RNA content was normalized to GAPDH and expression levels were quantified using 2^-ΔΔCT^ method from at least three independent experiments performed in triplicate.

**Table 2.** List of Taqman Probes for qPCR

| Probe | Product Number | Species |
| --- | --- | --- |
| Mig6 | Hs0021960_ml | Human |
| Chop | Hs00358796_gl | Human |
| Bip/Hsp5a | Hs00607129_gh | Human |
| Socs4 | Hs00328404_s1 | Human |
| Socs5 | Hs00751962_s1 | Human |
| Lrig1 | Hs00394267_m1 | Human |
| GAPDH | Hs02758991_g1 | Human |
| Mig6 | Rn01520745_m1 | Rat |
| Chop | Rn01458526_m1 | Rat |
| Bip/Hsp5a | Rn01435769_g1 | Rat |
| Socs4 | Rn01414734_m1 | Rat |
| Socs5 | Rn01769079_m1 | Rat |
| Lrig1 | Rn01421201_m1 | Rat |
| GAPDH | 4352338E | Rat |

### Mig6 overexpression and siRNA transfection

For gene overexpression studies, cells were transduced with adenoviral vectors driving the expression of green florescence protein (GFP) or Mig6 under the control of the cytomegalovirus (CMV) promoter for 24 h as previously described (46). Media was replaced with complete media the follow day and ligand stimulation was conducted 48 h after transduction. For RNA interference studies, cells were seeded at 50% confluency and forward transfected with siRNAs (30 pmol) using Lipofectamine RNAiMAX (Invitrogen, Carlsbad, CA) according to manufacturer’s guidelines. Sequences for RNAi are as follows: Mig6-1 forward 5’-CGAUAAUAGAACUAGUGACtt-3’ reverse 5’-GUCACUAGUUCUAUUAUGtt-3’, Mig6-2 forward 5’-GCUAUGUGUCUGACCAAAAtt-3’ reverse 5’-UUUUGGUCAGACAUAGCtg-3’, and the control *GL-2* siRNA forward 5’-CGUACGCGGAAUACUUCGAtt-3’ reverse 5’-UCGAAGUAUUCCGCGUACGtt-3’ as previously described (36). The following day the media was replaced with complete media. Ligand and drug treatments were conducted as previously described.

### Statistical analysis

All experiments were performed at least three independent times. Significance was determined by one- or two-tailed Student’s *t*-test or one- or two-way ANOVA with Bonferroni multiple comparison tests where appropriate. Differences were considered significant when *P* < 0.05. Data are reported as means ± SEM.

## RESULTS

### Palmitate and ER stress decreases EGFR activation and activates Mig6 expression

We previously determined that cellular stress induces Mig6, which impairs EGFR signaling and promotes apoptosis in pancreatic beta cells (6). To determine if EGFR activation and subsequent signaling are also inhibited by fatty acid-induced ER stress, primary rodent hepatocytes were treated with the saturated fatty acid palmitate prior to EGF stimulation. Indeed, palmitate treatment decreased ligand-mediated EGFR activation and prevented the induction of downstream ERK phosphorylation (Figure 1A-C). This signaling inhibition occurred in the context of increased intrinsic cell death as measured by caspase 3/7 activity (Figure 1D).

**Figure 1.**
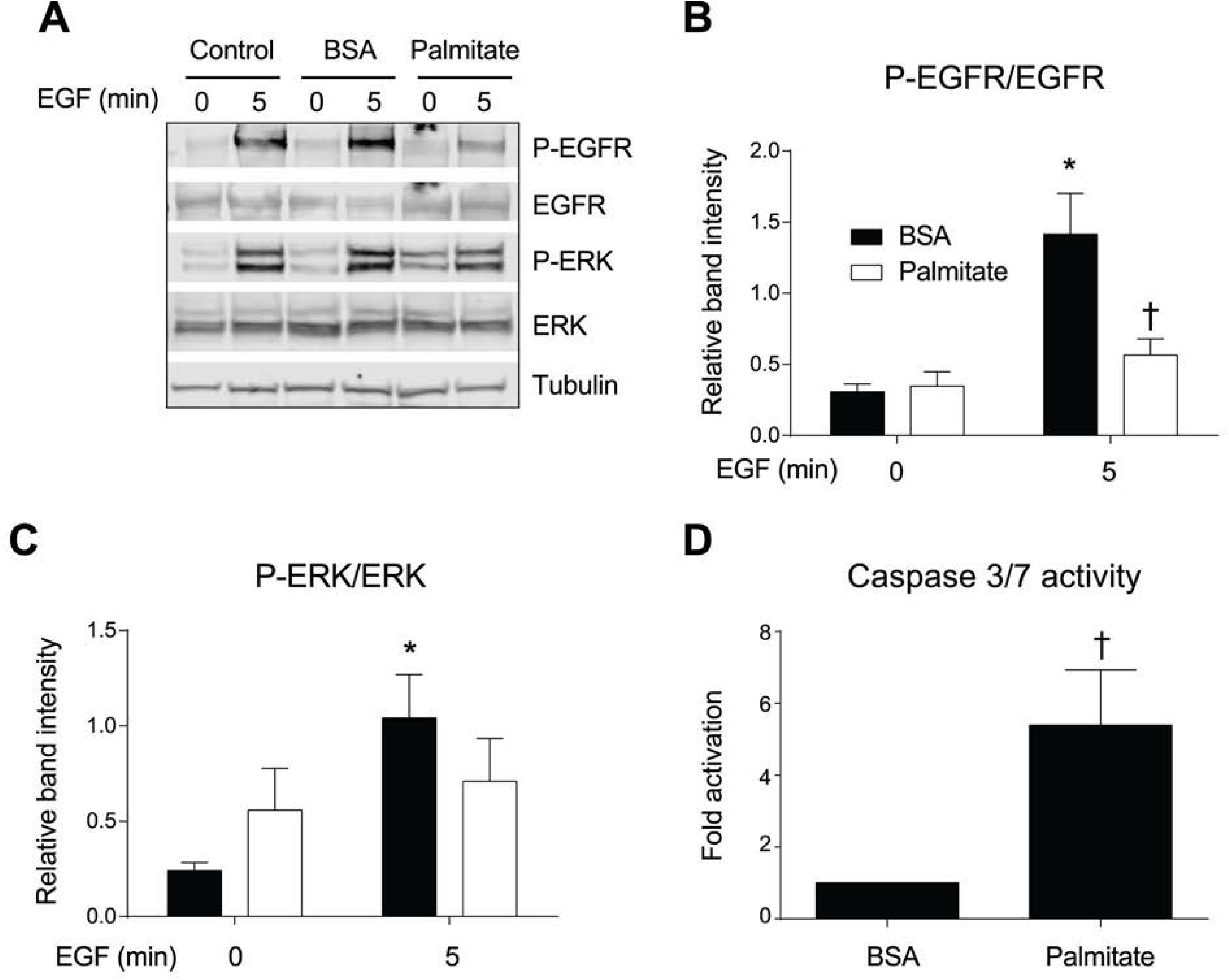
Palmitate decreases EGFR activation in primary hepatocytes and activates caspase 3/7. Primary rat hepatocytes were treated with BSA or palmitate (750 μM) for 48 h. 24 h prior to rrEGF (50ng/mL) media was replaced with serum free media. EGFR and ERK activation were assessed by Western blot. Shown is a representative blot (A). P-EGFR and P-ERK were normalized to EGFR and ERK, respectively (B,C). Caspase 3/7 activity was measured by luminescence following palmitate treatment for 48 h (D). Data are reported as means ± SEM; n = 3-4. *, p < 0.05 vs. untreated cells; †, p < 0.05 vs. BSA treated cells.

The magnitude and duration of EGFR signaling can be modulated by several inducible feedback inhibitors, including *Socs4, Socs5, Lrig1*, and *Mig6*. Thus, inhibitor expression levels were assessed in palmitate-treated H4IIE cells. Under these fatty acid-induced ER stress conditions, as noted by elevated levels of *Bip* and *Chop*, only *Mig6* gene expression was significantly increased whereas the other inhibitors were not induced (Figure 2).

**Figure 2.**
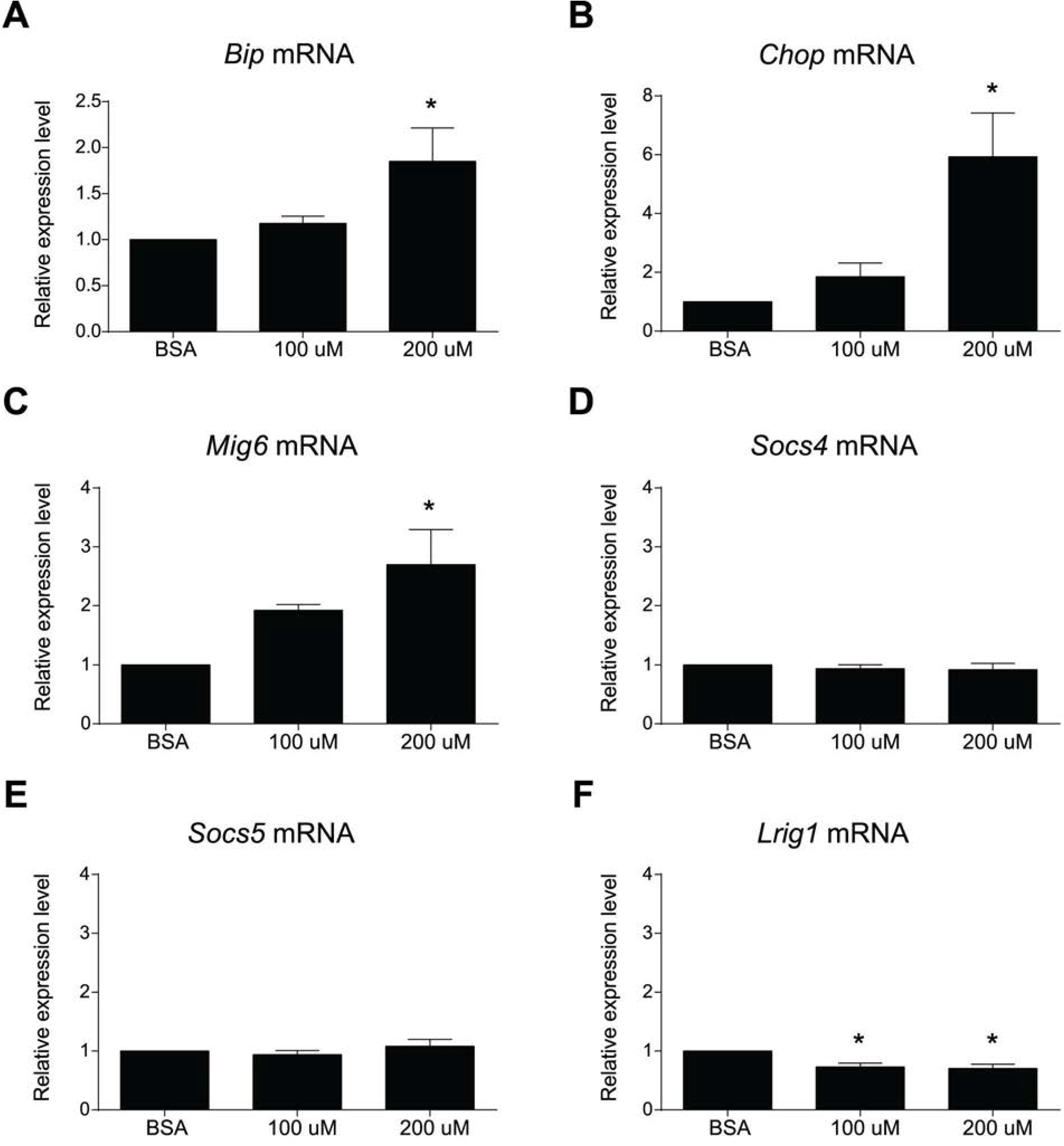
Palmitate-induced ER stress increases Mig6 expression in H4IIE cells. H4IIE cells were treated with vehicle (BSA) or palmitate for 8 h at indicated dose indicated in serum free media. mRNA expression was determined by qPCR. ER stress was confirmed by Bip and Chop mRNA expression (A,B). *Mig6, Lrig1, Socs4*, and *Socs5* mRNA were also determined (C-F). Data are reported as means ± SEM; n = 4. *, p < 0.05 vs. DMSO.

Diet-induced obesity triggers a variety of pathological responses in the liver including inflammation, insulin resistance, steatosis, and ER stress (18, 33). Because we have previously established that Mig6 is upregulated during ER stress in non-hepatic tissues (6), we sought to directly measure the effects of chemically-induced ER stress on EGFR inhibitors in hepatocytes, with the rationale being that ER stress in response to palmitate treatment promotes Mig6 induction. H4IIE cells were treated with the ER stress inducer thapsigargin, which inhibits the sarcoendoplasmic reticulum calcium transport ATPase, and ER stress was again confirmed by elevated *Bip* and *Chop* expression (Figure 3A,B). Like palmitate exposure, thapsigargin treatment increased *Mig6* gene expression in both dose-responsive and time-dependent manners (Figure 3C). Further, ER stress, as indicated by elevated *Bip* expression, increases *Mig6* transcript and protein levels in human HepG2 cells (Figure 4). Importantly, no changes were observed in *Socs4, Socs5*, and *Lrig1* gene expression in H4IIE during thapsigargin treatment compared with DMSO, and only *Socs4* gene expression was increased during thapsigargin treatment in HepG2 cells (data not shown). Thus, Mig6 alone appears to be the primary, or even exclusive, form of negative feedback inhibition of EGFR in hepatocytes during both fatty acid- and chemically-induced ER stress.

**Figure 3.**
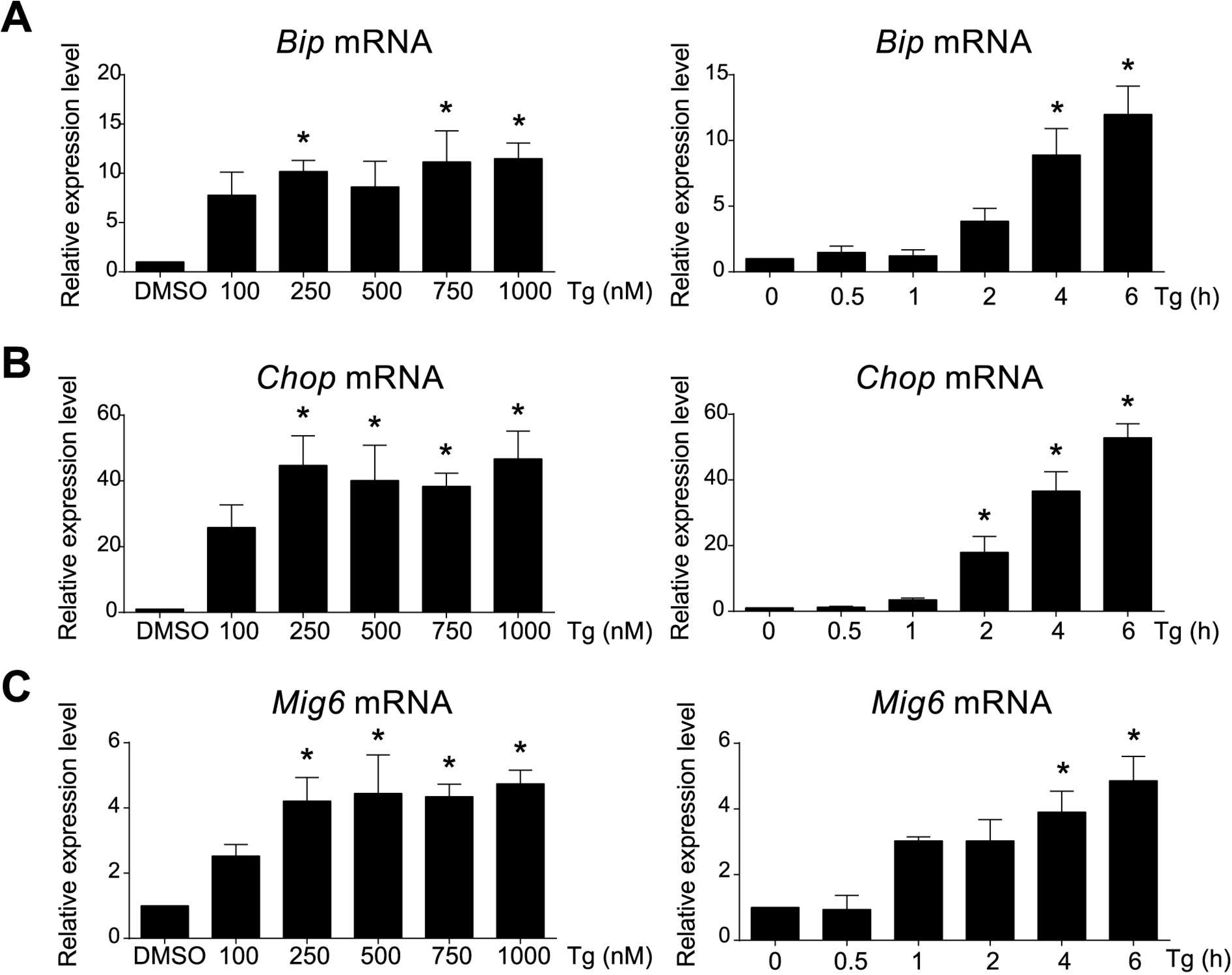
ER stress induces Mig6 in rat H4IIE cells. H4IIE cells were treated with vehicle (DMSO) or 1.0 μM thapsigargin (Tg) for 6 h or dose (nM) and time (h) indicated. *Mig6, Chop* and *Bip* mRNA (A-C) expression were assessed by qPCR. Data are reported as means ± SEM; n = 4. *, *p* < 0.05 vs. DMSO.

**Figure 4.**
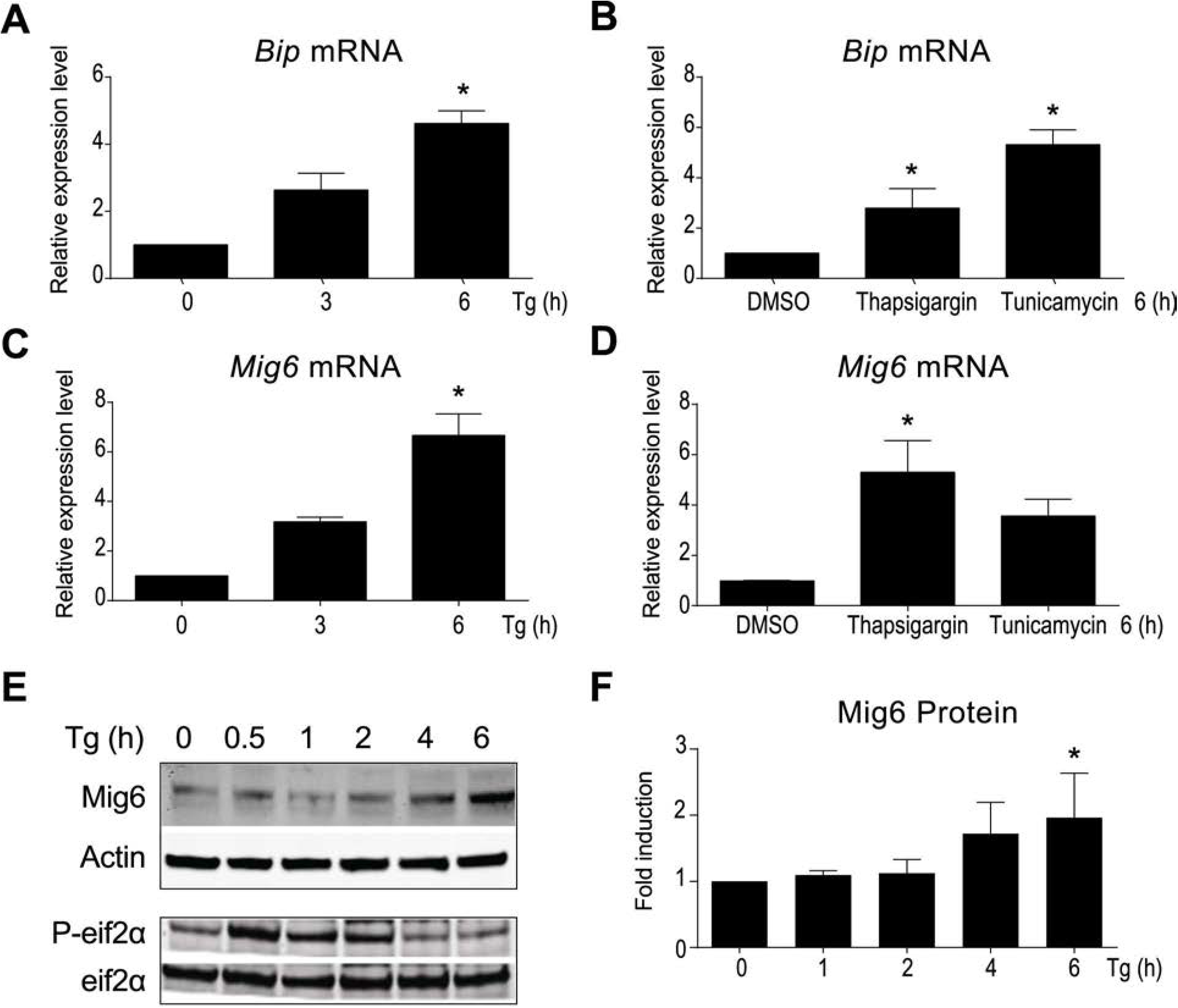
ER stress induces Mig6 in HepG2 cells. HepG2 cells were treated with vehicle (DMSO), 1.0 μM thapsigargin (Tg), or 1.0 μM tunicamycin for hours (h) indicated. *Bip* and *Mig6* mRNA (A-D) expression were assessed by qPCR. Mig6, P-eif2α, and eif2α protein expression were determined by Western blot. Shown is a representative blot (E). Mig6 was normalized to actin (F). Data are reported as means ± SEM; n = 4. *, p < 0.05 vs. DMSO.

### ER stress decreases EGFR activation and stabilizes Mig6 expression

In agreement with a previous report (46), adenoviral-mediated flag-tagged Mig6 overexpression is sufficient to decrease EGF-mediated EGFR activation as measured by phosphorylation of EGFR^Y1068^ at 5 min compared with controls (Figure 5A,B). Additionally, Mig6 overexpression reduced phosphorylation of the downstream kinase ERK1/2 (Figure 5A,C). Interestingly, under these conditions in both human and rat cell lines, EGF does not induce AKT^T308^ phosphorylation (data not shown). Given that maximal activation of the EGFR signaling cascade occurred at 5 min post-EGF stimulation, this time point was used in subsequent *in vitro* studies.

**Figure 5.**
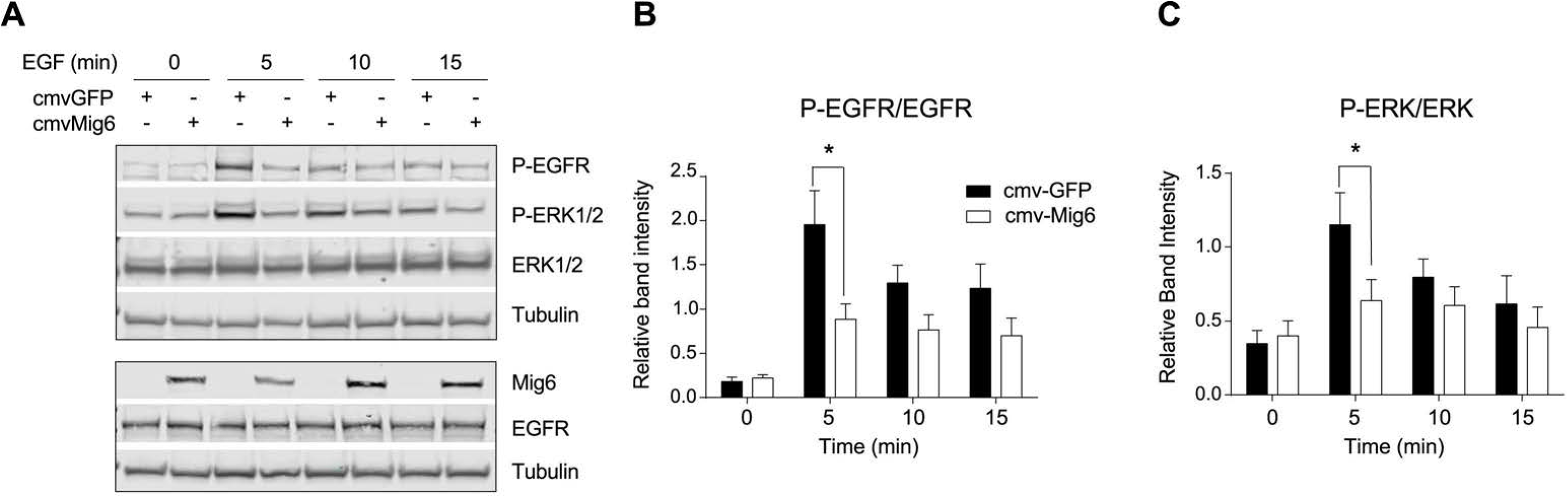
Mig6 overexpression reduces EGF-stimulated EGFR and ERK phosphorylation. HepG2 cells were transduced with control GFP- or Mig6-overexpressing adenoviruses. After 24 h media was replaced with serum free media and the following day cells were treated with vehicle or rrEGF (50 ng/mL) for times indicated. Shown is representative Western blot (A). p-EGFR and EGFR were normalized to tubulin and P-ERK was normalized to ERK protein then quantified (B,C). Data are expressed as means ± SEM; n = 4. *, *p* < 0.05 vs. cmv-GFP.

As Mig6 is induced during ER stress and Mig6 overexpression decreases EGFR activity, we hypothesized that EGFR activity is decreased during chemically-induced ER stress. Again, we used thapsigargin to induce hepatic ER stress in HepG2 cells (Figure 6). During this stress condition, EGF-mediated EGFR activation is markedly attenuated and apoptosis is activated, as indicated by the presence of activated, cleaved caspase 3 on the immunoblot (Figure 6A-C). Under similar ER stress conditions, we used the flag-tagged Mig6 adenovirus to demonstrate Mig6 stability was markedly increased, indicating Mig6 is actually stabilized during ER stress (Figure 6D-E).

**Figure 6.**
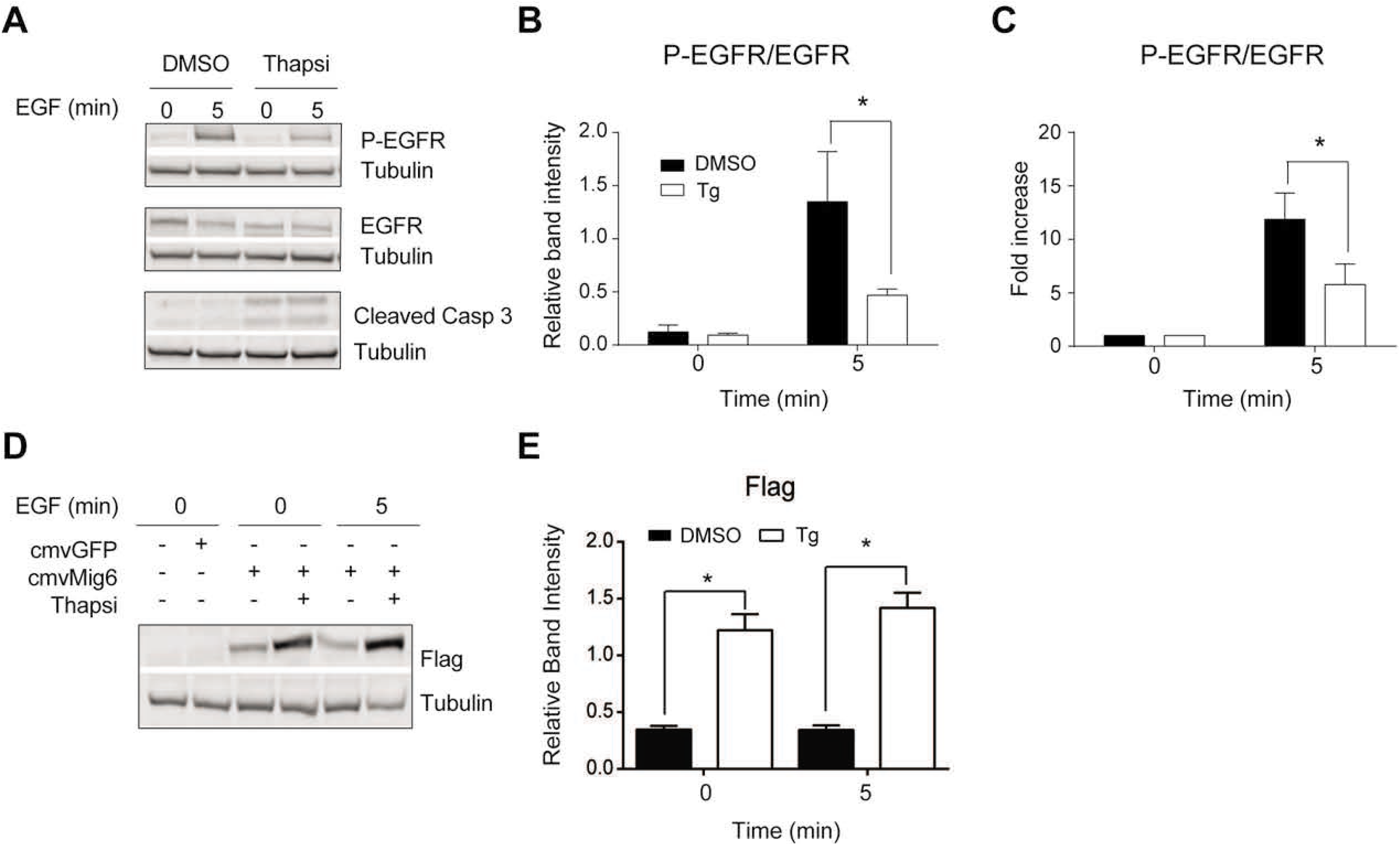
Thapsigargin decreases EGFR activation, induces caspase 3 activation, and stabilizes Mig6 protein expression. HepG2 cells were treated with vehicle (DMSO) or thapsigargin (1 μM) for 24 h in serum free media prior to rrEGF (50ng/mL) stimulation as indicated (A-C). HepG2 cells were transduced with control GFP- or Mig6-overexpressing adenoviruses. After 24 h media was replaced with serum free media and treated as above (D,E). EGFR activation, caspase 3 cleavage, and flag expression were assessed by Western blot. Shown are representative blots (A&D). P-EGFR and EGFR were normalized to tubulin then quantified (B, C). Flag-tagged Mig6 expression was normalized to tubulin and quantified (E). Data are reported as means ± SEM; n = 3 (A-C) and n =4 (D,E) *, *p* < 0.05 vs. DMSO.

### Mig6 knockdown enhances EGFR activation and reduces caspase 3/7 activity during ER stress

To establish that the suppression of EGFR activation during ER stress is indeed mediated by Mig6, HepG2 cells were pre-transfected with siRNAs targeting Mig6 (siMig6) or a scrambled control siRNA (siCon) and then exposed to thapsigargin (or vehicle) prior to ligand stimulation (Figure 7). Compared to siCon-transfected cells, siMig6-transfected cells have reduced Mig6 expression and increased EGFR phosphorylation during both control (i.e., vehicle-treated) and ER stress conditions (Figure 7A,B). This observation confirms Mig6 directly inhibits EGFR activation during ER stress.

**Figure 7.**
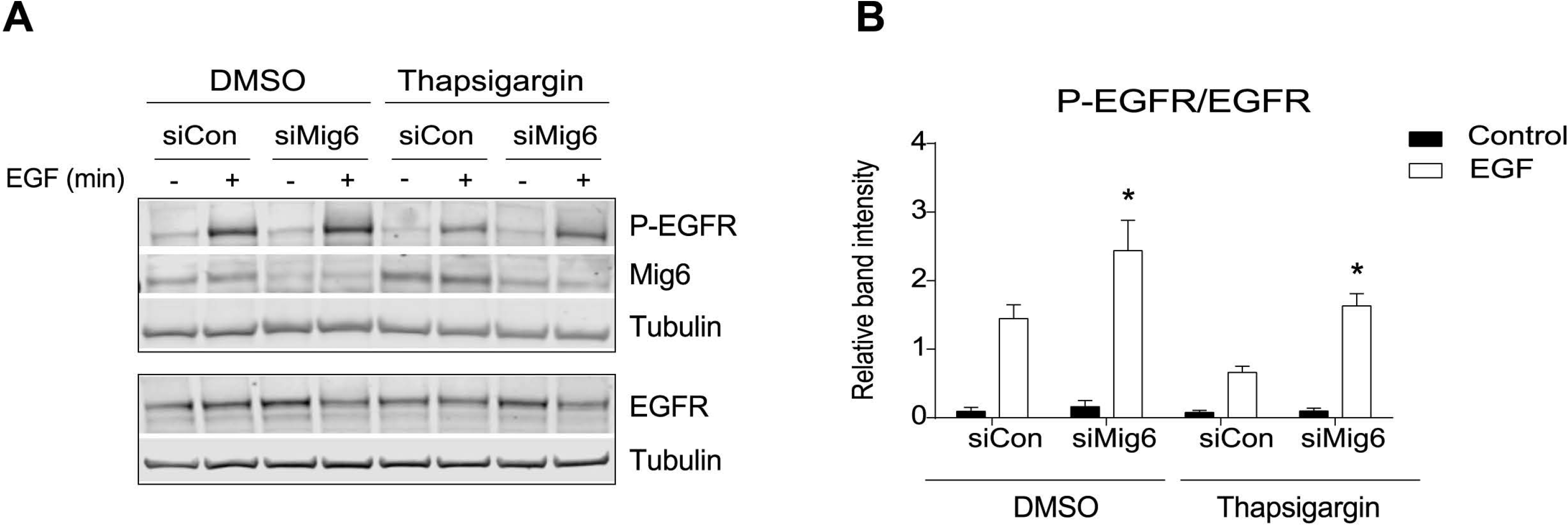
Mig6 ablation rescues EGFR activation during ER stress. HepG2 cells were pretreated with siRNAs directed against control (siCon), or Mig6 (siMig6) for 48 h prior to ligand stimulation. Media was replaced with serum free media and either vehicle (DMSO) or thapsigargin (1.0 μM) 16 h prior to EGF stimulation (50ng/mL for 5 min). EGFR activation and Mig6 expression were determined by Western blot. A representative blot is shown (A). P-EGFR was normalized to total EGFR (B). Data are reported as means ± SEM; n = 3. * *p* < 0.05 vs. siControl.

To determine the functional significance of decreased EGFR signaling during ER stress, HepG2 cells were treated with palmitate following siRNA-mediated suppression of Mig6. Palmitate induced both ER stress and Mig6 transcript expression (Figure 8A,B). Mig6 suppression reduced caspase 3/7 activity during palmitate treatment compared to siControl-transfected cells (Figure 8C). These data indicate dampening the inhibitory actions of Mig6 via siRNA restores EGFR signaling, leading to reduced caspase 3/7 activation and enhanced survival in HepG2 cells during ER stress.

**Figure 8.**
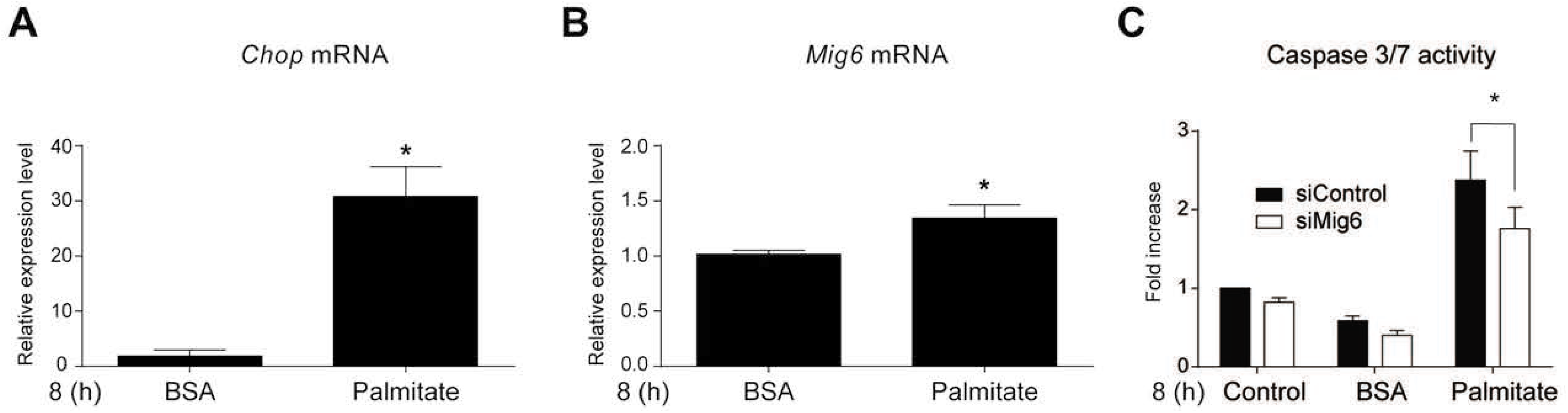
Mig6 knockdown during palmitate-induced ER stress decreases caspase 3/7 activity in HepG2 cells. HepG2 cells were incubated in serum free media containing either BSA or palmitate (750 uM) for 8 h and. For caspase 3/7 activity cells were treated with nothing or oligos directed against control or Mig6 for 48 h. Following transfection, media was replaced and cells were treated as before. Caspase 3/7 activity was measured using chemiluminescence. *Chop* and *Mig6* mRNA were measured by qPCR (A,B) and caspase 3/7 activity is expressed as fold increased luminescence from control non-treated cells (C). Data are reported as means ± SEM; n=3 *p* < 0.05 vs. palmitate treated siControl.

### Mig6 protein expression is elevated during obesity

Because diet-induced obesity drives hepatic ER stress, we sought to determine the impact of diet-induced obesity on hepatic Mig6 expression. Male C57Bl/6J mice were fed diets containing either 15 or 60% kcals from fat (low vs. high fat diets, LFD vs. HFD, respectively) for 15 weeks. As expected, HFD-fed mice had increased body weight and pre-diabetic hyperglycemia and hyperinsulinemia compared to LFD-fed controls (Figure 9A-C). Importantly, hepatic Mig6 protein expression but interestingly not mRNA expression was increased in HFD-fed animals compared to LFD-fed controls (Figure 9D-F), suggesting Mig6 protein stability might be increased during obesity.

**Figure 9.**
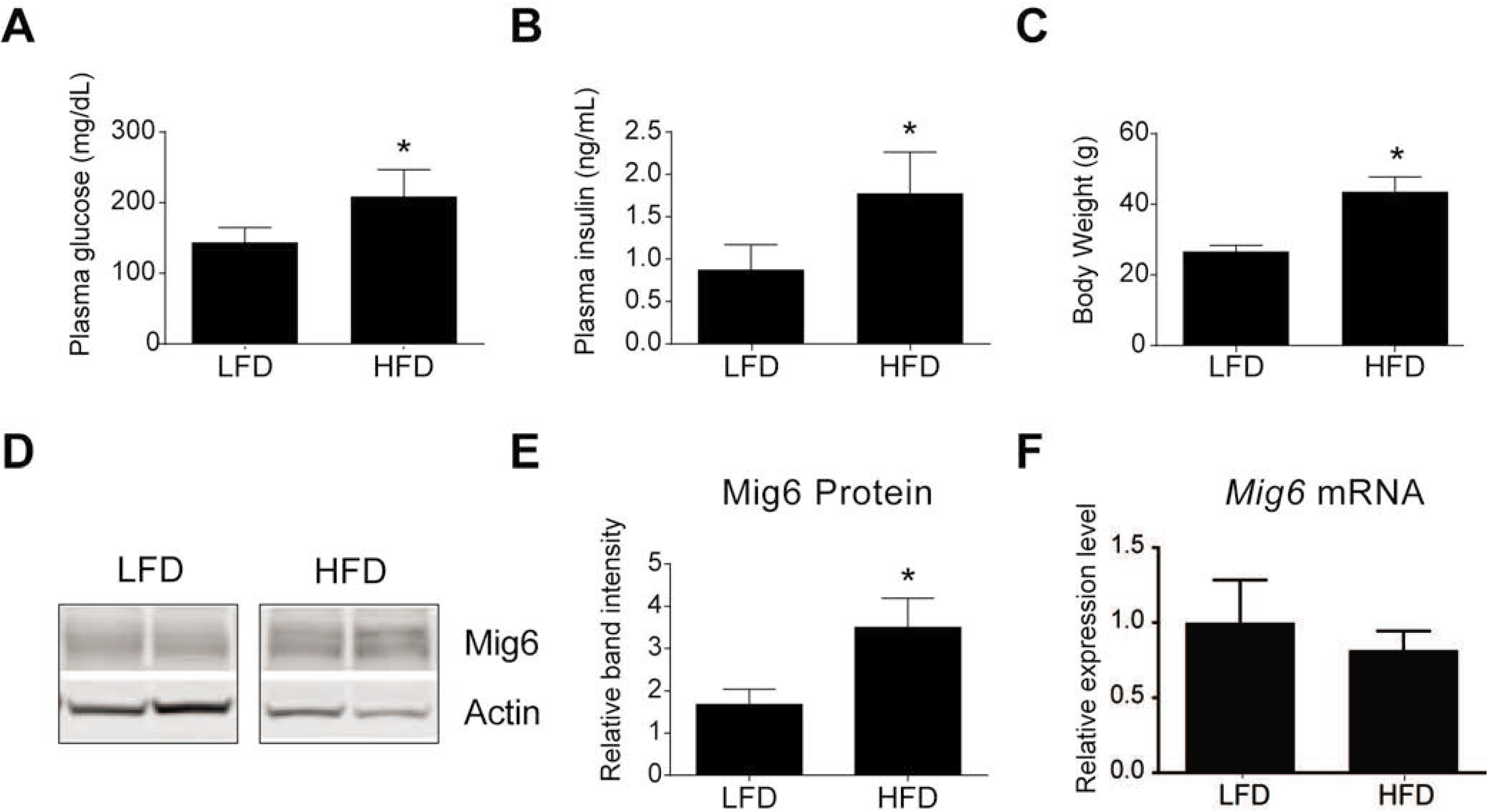
Mig6 in increased in diet-induced obesity and prediabetes. 8 week old male C57Bl/6 mice (n=4) were fed a LFD (15% calories from fat) vs. HFD (60% calories from fat) for 15 weeks. HFD mice develop higher blood glucose (A), plasma insulin levels (B), and body weight (C) compared to LFD mice. Representative Western blots of liver tissue for Mig6 protein (D) and quantifications normalized to actin (E). *Mig6* mRNA was measured by qPCR (F). Data are reported as means ± SEM; n = 4. *, *p* < 0.05 vs. LFD.

## DISCUSSION

Hyperlipidemia from over-nutrition and obesity increases hepatic lipid uptake and deposition (i.e., steatosis). Clinically, steatosis is often considered benign, requiring a “second hit” to initiate the necroinflammation and steatohepatitis-like lesions responsible for reduced patient survival during the transition from simple steatosis to non-alcoholic steatohepatitis (NASH) and cirrhosis (2, 40). Hepatic necroinflammation and steatohepatitis provoke oxidative stress-induced DNA damage and caspase activation (13, 43). In response to these insults, hepatocytes and local immune cell populations trigger potent activators of hepatic DNA damage repair and synthesis, including EGFR, c-Met/HGF receptor, and TGF-β receptor and their ligands. These mitogens and their cognate receptors have been extensively investigated in healthy livers (12, 29, 30). However, the molecular mechanisms responsible for their inhibition during obesity- and ER stress, which compromises cell survival, have remained largely unexplored.

One possible explanation for compromised hepatic survival in obesity could be stress-mediated feedback inhibition of RTKs. Here, we have established that a negative feedback inhibitor of EGFR, Mig6, is upregulated during diet-induced obesity, hyperglycemia, and hyperinsulinemia in mice. Because obesity and impaired insulin signaling cause hepatic ER stress, we also employed ER stressors to inhibit EGFR signaling and induce Mig6 expression in cultured hepatocytes. The increased Mig6 expression was necessary for inhibiting EGFR activation, as evidenced by rescued EGFR activation following Mig6 suppression during stress conditions. Although Mig6 gene expression was not elevated in livers of obese mice, we speculated Mig6 protein stability may be enhanced based on our *in vitro* studies (Figure 6). Importantly, Mig6 is sufficiently upregulated and Mig6 mRNA and protein stability are enhanced during ER stress (here and 26). We have previously demonstrated Mig6 transcripts escape translational blockade during ER stress conditions in β-cells (7).

Similar to diet-induced EGFR inactivation, previous studies have revealed hepatic steatosis decreased EGFR expression and activation in genetically obese, *ob/ob* (i.e., leptin-deficient) mice (10, 17). Although the exact mechanisms remain elusive, EGFR expression can be restored with growth hormone administration in *ob/ob* mice (10). Moreover, leptin administration to *ob/ob* mice fails to rescue the regeneration response following partial hepatectomy, suggesting that fatty-liver pathologies might dampen the regenerative response through EGFR inactivation (25). More importantly, these studies failed to explore the potential contribution from endogenous feedback inhibitors such as Mig6, which we have now established to be elevated during diet-induced obesity.

EGFR activation is tightly controlled by both receptor-mediated endocytosis and ligand-activated feedback inhibition (39). Though several endogenous inhibitors exist, few have been examined during cellular stress conditions such as ER stress. Here, we demonstrate the largely exclusive upregulation of Mig6 during ER stress in liver. Additionally, there was a small but significant decrease in *Lrig1* mRNA expression during thapsigargin-induced ER stress in rat hepatoma cells. Unlike LRIG1, SOCS4, and SOCS5, Mig6 inhibits both receptor activation and enhances receptor degradation (39). Therefore, expression of EGFR and its ligands alone may not be capable of enhancing proliferation and survival during the mitogenic phase of regeneration if its activation is impaired by Mig6 (38, 39). During feedback inhibition, Mig6 binds to the catalytic domain of EGFR through its ERBB binding domain, acting as a docking site for endocytic proteins (48, 50). ER stress may potentially facilitate this binding interaction. However, increased binding of Mig6 to EGFR during stress remains to be demonstrated.

Obesity-induced ER stress causes decreased regeneration following partial hepatectomy (41). Conversely, administration of the chemical chaperone tauroursodeoxycholic acid alleviates ER stress and restores hepatic regeneration in rats (31). Given that ER stress induces Mig6 and Mig6 halts cell cycle progression from G1 to S phase, Mig6 could potentially inhibit proliferation during ER stress. For example, proliferation requires massive ER expansion and protein translation. If sufficient stress on the ER persists, intrinsic mitogen inhibitors may become activated to halt proliferation until stress is resolved or the cell undergoes regulated apoptosis. Mig6 expression enhances apoptosis in several cell types (38, 26, 36, 47, 6). Importantly, active proliferation is observed in both non-fulminant and fulminant hepatitis, indicating proliferative response *alone* is not sufficient to prevent cell death (22, 44). Here, we have demonstrated Mig6 knockdown reduces FFA-induced apoptosis through decreased caspase 3/7 activity. As previously discussed, hepatic lipid storage increases during obesity and the FFAs entering the liver are esterified and stored as triglycerides. However, a subset of lipids become diacylglycerols (DAG) which activate atypical protein kinase C’s (PKC) (37). Interestingly, Mig6 is induced by DAG analogues phorbol esters, possibly suggesting additional induction mechanisms during FFA stress (15). In addition to DAG, ceramides increase during FFA uptake, though Wei et al. show palmitate induces ER stress and apoptosis independently of ceramides in liver cells (42).

Our ultimate goal is to elucidate the extent to which Mig6 mediates the impaired hepatic regeneration during obesity and if this occurs through enhanced cell death, decreased proliferation, or both. Mig6 could potentially halt the proliferative response of a steatotic liver following a hepatectomy. Mig6 ablation permits enhanced hepatic regeneration through heightened EGFR activation and signaling (36). Although these studies were not conducted in the context of obesity compromised livers and Mig6 whole body knockout have poor viability (4, 14, 20, 51). Additionally, under normal conditions, liver-specific deletion of Mig6 leads to hepatomegaly, without any reports of hepatic dysfunction or hepatocellular carcinoma development, suggesting Mig6 restricts liver proliferation (24). Physiologically, EGFR controls the initial stages of regeneration when hepatocytes are in the first round of the cell cycle. EGFR activation leads to downstream kinase cascades including ERK1/2 activation and subsequent induction of cyclins, allowing the transition between G1 and S phase of the cell cycle (9, 32). Thus, Mig6 inhibits the G1 to S phase transition (11). Although the regenerative response is subjected to several mitogen receptors, deletion of individual receptors, or their ligands, only delays or impairs regeneration without complete abrogation. Challenged by obesity-induced ER stress, the liver loses the ability to fully regenerate following partial hepatectomy, suggesting further mechanisms remain.

Here, we describe a mechanism where EGFR is inhibited during ER stress, and we speculate that other pro-survival signaling axes like HGF/c-met may also be compromised during ER stress. We hypothesize unrestrained EGFR signaling in the context of impaired repair and proliferation (e.g., impaired regeneration during diet-induced obesity) may accelerate growth and recovery of otherwise impaired tissue. Future studies using mice lacking Mig6 in the liver may provide further insight into the complete impact of EGFR signaling inhibition during regeneration, survival, and repair.

## ACKNOWLEDGEMENTS

We would like to thank Scott Colvin, MS and Dr. Angelina Hernandez for thoughtful discussions. We would also like to thank the Indiana University Center for Diabetes and Metabolic Diseases as well as Dr. David Morris for providing invaluable samples and resources.

